# Trafficking of Connexin36 (Cx36) in the early secretory pathway

**DOI:** 10.1101/2024.03.25.586643

**Authors:** Stephan Tetenborg, Fatemeh Ariakia, Elizabeth Martinez-Soler, Eyad Shihabeddin, Ignacio Cebrian Lazart, Adam C. Miller, John O’Brien

**Affiliations:** College of Optometry, University of Houston, Houston, TX, USA; Institute of Neuroscience, Department of Biology, University of Oregon, Eugene, OR, USA; MD Anderson Cancer Center UTHealth Graduate School of Biomedical Sciences, Houston, TX, USA; Instituto de Histología y Embriología de Mendoza (IHEM)-CONICET, Facultad de Ciencias Médicas, Universidad Nacional de Cuyo, Mendoza 5500, Argentina; Centre for Genomic Regulation (CRG), The Barcelona Institute for Science and Technology, Dr. Aiguader 88, Barcelona 08003, Spain

## Abstract

Gap junctions formed by the major neuronal connexin Cx36 function as electrical synapses in the nervous system and provide unique functions such as synchronizing activities or network oscillations. Although the physiological significance of electrical synapses for neuronal networks is well established, little is known about the pathways that regulate the transport of its main component: Cx36. Here we have used HEK293T cells as an expression system in combination with siRNA and BioID screens to study the transition of Cx36 from the ER to the cis Golgi. Our data indicate that the C-terminal tip of Cx36 is a key factor in this process, mediating binding interactions with two distinct components in the early secretory pathway: the COPII complex and the Golgi stacking protein Grasp55. The C-terminal amino acid valine serves as an ER export signal to recruit COPII cargo receptors Sec24A/B/C at ER exit sites, whereas the PDZ binding motif “SAYV” mediates an interaction with Grasp55. These two interactions have opposing effects in their respective compartments. While Sec24 subunits carry Cx36 out of the ER, Grasp55 stabilizes Cx36 in the Golgi as shown in over expression experiments. These early regulatory steps of Cx36 are expected to be essential for the formation, function, regulation and plasticity of electrical synapses in the developing and mature nervous system.

## Introduction

In the nervous system neurotransmission occurs in two different modalities: either via chemical or electrical synapses. Chemical synapses rely on the secretion of a neurotransmitter by a presynaptic neuron and the transformation of the corresponding signal into a change of the post synaptic membrane potential (1). Neurotransmission at electrical synapses, on the contrary, is instantaneous and mediated via intercellular channels that allow the bidirectional flow of ionic currents between connecting neurons (2). Given the fast properties of electrical transmission, it can synchronize neuronal populations, which is a key factor in a variety of different brain functions, including hormone secretion, generation of network oscillations and the coordination of motor functions (3, 4).

Structurally, electrical synapses are defined as gap junctions, which are made of dense clusters of tens to thousands of intercellular channels forming direct cytoplasmic connections (5). The molecular building blocks of these channels are connexins, a family of membrane proteins that arose during the early chordate evolution (6). A characteristic feature of connexins, is their ability to oligomerize into hexameric hemichannels (also called connexons). Depending on the connexin isoform, these channels are either assembled in the endoplasmic reticulum (ER) or the Golgi apparatus, and transported to the gap junction where they dock to opposing connexons from the neighboring cell to form a functional intercellular channel (7, 8).

The connexin gene family consists of 21 different isoforms in the human genome with each variant sharing a conserved protein structure including four transmembrane domains, two extracellular loops and three intracellular domains (9). Among all connexin isoforms in mammals, connexin 36 (Cx36) is often referred to as “the major neuronal connexin” because of its widespread expression in neurons (10). Gap junction channels formed by Cx36 exhibit an unusually low single channel conductance but a tremendous degree of plasticity, enabling electrical synapses to modify the strength of transmission (11–14). While little is known about the molecular mechanisms that regulate the transport of Cx36 in neurons, it is evident that the PDZ binding motif (PBM) and its interaction with the scaffold protein ZO-1 are critical for electrical synapse formation (15). The significance of this interaction has been illustrated in a variety of examples. For instance, the expression of a Cx36-EGFP fusion protein in a Cx36 knock-out mouse background has been shown to interfere with gap junction formation in retinal neurons and other brain regions such as the cerebellum or the olfactory bulb. These deficits are thought to arise because the C-terminal GFP tag prevents access to the PBM in Cx36, resulting in a loss of association with important binding partners such as ZO-1 (15). Interestingly, no defects in synapse formation are observed when Cx36-EGFP is co-expressed alongside wild type Cx36. This rescue effect can be explained by connexon hexamers being constructed of both wildtype Cx36 and Cx36-EGFP, creating chimeric hemichannels with sufficient functional C-terminal PBMs to compensate for the loss of a functional PBM in Cx36-EGFP (15, 16). In zebrafish Mauthner cells it was demonstrated that genetic ablation of the ZO-1 homolog prevents electrical synapse formation, leading to a failure in electrical synaptic transmission (17). Consistent with these findings, Flores et al. (18) described that intradendritic application of peptides mimicking a C-terminal portion of the Cx36 homolog into goldfish Mauthner cells reduced electrical synaptic transmission. These studies have suggested that PDZ binding interactions between Cx36 and ZO-1 (and their homologues) occur at the level of the plasma membrane to recruit or stabilize intercellular gap junction channels. Yet, PBM-mediated interactions could regulate other steps of Cx36 hemichannel assembly earlier in the secretory pathway. Indeed, previous work from our group found that the C-terminal tip of Cx36 functions as an ER export signal, as Cx36 mutants that lacked the PBM displayed severe ER retention resulting in the formation of ER derived removal vesicles (19). In line with this observation, we identified perinuclear accumulations of Cx36-EGFP in AII amacrine cell somas of the Cx36-EGFP/Cx36 KO mouse strain, resembling the ER retention phenotype that was observed in transiently transfected cells (19). Thus, the loss of gap junction formation described for Cx36-EGFP/Cx36 KO mice (15, 16) may not be solely attributable to a lack of association with ZO-1 but could reflect a secretory deficit.

In the present study we have dissected the underlying mechanisms guiding the transport of Cx36 in the early secretory pathway. By combining BioID and RNA interference, we find the Cx36 C-terminus drives distinct interactions with the COPII complex at ER exit sites and with Grasp55 in the cis Golgi. At the ER, the C-terminal amino acids tyrosine and valine function as an export signal recruiting the COPII cargo receptors Sec24A and B at exit sites which target Cx36 into export vesicles. After vesicles transition to the Golgi, Cx36 binds to the scaffold protein Grasp55 via the PBM, likely concentrating the connexin in the Golgi. In summary, our study provides a first description of the events that regulate the transition of a neuronal gap junction protein from the ER into the Golgi.

## Material and methods

### Cell Culture and Transient Transfection

Human embryonic kidney 293T cells (HEK293T/17; catalog #CRL-11268; ATCC, Manassas, VA, USA; RRID: CVCL_1926) and Grasp55 knockout HEK293T cells ((20), generous gift of Dr. Jayanta Debnath, University of California at San Francisco) were grown in Dulbecco’s Modified Eagle Medium (DMEM) supplemented with 10% fetal bovine serum (FBS), 1% penicillin and streptomycin, and 1% non-essential amino acids (all Thermo Fisher Scientific, Rockford, IL, USA) at 37°C in a humidified atmosphere with 5% CO2. For immunostainings 4.5 x 10^5^ cells were grown in 35mm dishes with poly-l-lysine (0.01 % for 30 min at RT) coated coverslips inside. Cells were transfected at the next day with 1µg DNA and 5µl Geneporter 2 (Genlantis, San Diego, CA). For Immunoprecipitations 1.5 x 10^6^ cells were plated onto 60 mm dishes and transfected at the next day with 4µg DNA and 10 µl Geneporter 2.

### RNA interference

For knock-down studies 1.66×10^5^ HEK293T cells were plated into 35 mm dishes. At the next day cells were transfected with 100nM siRNA using 5µl lipofectamine^TM^ RNAiMAX (ThermoFisher) in DMEM containing 10% FBS. 48h after transfection siRNA treated cells were co-transfected with 333ng of Cx36 pcDNA and 100nM siRNA. The following siRNA sequences were used as duplexed RNA in this study. Random seq1: 5‘GCAAUGAGCGUGUGCGACCUAUTT3‘, Random seq2: 5‘GCAAUGAGCGUGUGCGACCUAUTT3‘, Sec24A seq1:<colcnt=3>

5‘UGUAUCCUUGGCUCACUGACUCTT3‘ Sec24A seq2:

5‘UGUAUCCUUGGCUCACUGACUCTT3‘, Sec24B seq1:

5‘AUAUACGUUCGACAGGAUCGGCTT3‘, Sec24B seq2:

5‘GCCGAUCCUGUCGAACGUAUAUTT3‘, Sec24C seq1:

5‘GCUUCUAGUAUCCAGACUCTT3‘. Sec24C seq2: 5‘GAGUCUGGAUACUAGAAGCTT3‘ Sec24D seq1: 5‘AGCCUCCUGGGUAAGACAGUUGTT3‘, Sec24D seq2: 5‘

5‘CAACUGUCUUACCCAGGAGGCUTT3‘. Syntaxin5 sense:

5‘CAAUAGCCUCAACAAACAAAUUTT3‘ Syntaxin5 anti-sense: 5‘AAUUUGUUUGUUGAGGCUAUUGTT3‘, GOPC seq1:

5‘UUUUCUAAUUGGACCAACACCUUGTT3‘, GOPC seq2:

5‘CAAGGUGUUGGUCCAAUUAGAAAATT3‘. 3 different RNAi duplexes were used for knock-down of Ergic3: duplex1 seq1: 5‘AUUGAUUGAUUGUGAUAGUAAAAAA3‘, Ergic3<colcnt=3>

duplex1 seq2: 5‘UUUUUUACUAUCACAAUCAAUCAAUAG3‘; Ergic3 duplex2 seq1:

5‘AAGAUCAACAUCGAUGUACUUUUTC3‘, Ergic3 duplex2 seq2:

5‘GAAAAAGUACAUCGAUGUUGAUCUUCA3‘; Ergic3 duplex3 seq1:

5‘AAGAACCCAGAUACUAUUGAGCAGT3‘, Ergic3 duplex3 seq2:

5‘ACUGCUCAAUAGUAUCUGGGUUCUU-GA3‘.

### DNA constructs

Cx36 and Cx36S318ter were previously used in Tetenborg et al., (19). C-terminal Cx36 mutants were generated using the Q5 site directed mutagenesis kit (E0554S; New England Biolabs, Ipswitch, MA) and corresponding primer pairs introducing gaps into the Cx36 coding sequence. Grasp55-GFP (Addgene: 137708), Sar1(T39N)-YFP (Addgene: 128155), EGFP-Sec23A (Addgene: 666009), GFP-Sec61B (Addgene: 121159) were ordered from Addgene. Grasp55-GFP/G2A and Grasp55-GFPΔPDZ1 +2 were generated using site directed mutagenesis. The Venus-PDZ1 (ZO-1) was generated by site directed mutagenesis of the Venus-ZO-1 expression vector (Addgene: 56394) using primer pairs that delete the entire ZO-1 coding sequence except for the first PDZ1 domain. V5-dGBP-TurboID was cloned into a mammalian expression vector and used previously (19, 21)

### Immunocytochemistry

Immunocytochemistry was carried as previously described (19). HEK293T cells were rinsed in PBS and fixed with 2% PFA in PBS for 15 min at room temperature. The coverslips were washed 3 x 10 min with PBS and incubated in the primary antibody diluted in 10% normal donkey serum in 0.5% Tx-100 (PBS) overnight at 4°C. Primary antibodies used were Cx36 1:500 (MAB3045; Millipore, Burlington, MA, RRID:AB_94632), Sec22b 1:250 (186004; Synaptic Systems, Goettingen, Germany), Syntaxin5 1:500 (110 053; Synaptic Systems), Sec24A 1:250 (PA5-66043; ThermoFisher, RRID:AB_2662296), Sec24B 1:250 (12042; Cell Signaling Technology, RRID:AB_2797807), Sec24C 1:250 (PA5-59101; ThermoFisher, RRID:AB_2647077), Golgin97 1:250 (PA5-84015; ThermoFisher, RRID:AB_2791167), GM130 1:250 (12480; Cell Signaling Technology, RRID:AB_2797933), PTPN13 1:250 (PA5-72906; Thermofisher, AB_2718760), Grasp55 1:250 (MA5-24642; ThermoFisher, RRID:AB_2637257) and Grasp55 1:500 (10598-1-AP; Proteintech, RRID:AB_2113473). At the next day the coverslips were washed 3 times with PBS and incubated with the secondary antibodies (from donkey, 1:500, conjugated with Cy3, Alexa488, Alexa568, or Alexa647; Jackson Immunoresearch, West Grove, PA) diluted in 10% normal donkey serum in 0.5% Tx-100 (PBS) for 1h under light protected conditions at room temperature. The coverslips were mounted in Vectashield containing DAPI (Vector Laboratories, Burlingame, CA) and sealed with nail polish.

### Proximity ligation assay

The proximity ligation was assay was carried out using the Sigma duolink proximity ligation assay (Sigma, DUO92101) according to the manufacturer‘s instructions. Fixed HEK293T cells were permeabilized for 1h with 0.5 % Tx-100 in PBS. Afterwards cells were incubated in blocking solution for 1h at 37 °C and incubated in the primary antibody (Cx36 1:2000, Chemicon, MAB3045; Sec24B 1:1000; Cell Signaling; Grasp55 1:2000, Proteintech, 10598-1-AP) diluted in the supplied antibody diluent overnight at 4°C. At the next day coverslips were washed twice for 5 min in wash buffer A and incubated in the PLUS and MINUS PLA probes (each diluted 1:40 in dilution buffer) for 1h at 37°C. The coverslips were washed twice in wash buffer for 5 min and incubated in the ligation mix (ligase diluted 1:40 in corresponding buffer solution) for 30 min at 37°C. Afterwards coverslips were washed again in wash buffer A and incubated in the amplification mix (Polymerase diluted 1:80 in amplification buffer) for 100 min at 37 °C. Afterwards coverslips were washed 2×10 min in wash buffer B, once in 0.01x wash buffer B for 1 min and mounted with supplied mounting medium.

### BioID

BioID experiments were carried out as previously described (19). For BioID experiments 4×10^6^ HEK293T cells were plated onto 10 cm dishes. At the next day cells were co-transfected with 4 µg Cx36-EGFP, 4 µg Cx36 and 4 µg V5-dGBP-TurboID using 50 µl Geneporter 2 (Genlantis). 24 h after transfection HEK293T cells were treated with 50 µM Biotin in 10% DMEM for 3 h to induce biotinylation of proximal proteins. Afterwards, HEK293T cells were rinsed with PBS and transferred to a reaction tube on ice. Cells were lysed in lysis buffer (1% Triton X-100, 50 mM Tris pH 7.5, 250 mM NaCl, 5 mM EDTA, 50 mM NaF, 1 mM Na3VO4, 1 mM DTT and protease inhibitors (Roche) and sonicated 30 min. The cell lysates were centrifuged at 16,000g for 10 min. 10 mg of the supernatant was mixed with 275 µl of streptavidin beads myOne C1 (Thermofisher, Dynabeads MyOne Streptavidin C1) and incubated overnight at 4**°**C on a rotating platform. At the next day streptavidin beads were collected using a magnetic stand and washed in the following washing buffers (10 min for each washing step): 2x in wash buffer 1 (2% SDS), once with wash (0.1% deoxycholate, 1% Triton X-100, 500 mM NaCl, 1 mM EDTA and 50 mM Hepes, pH 7.5), once with wash buffer 3 (250 mM LiCl, 0.5% NP-40, 0.5% deoxycholate, 1 mM EDTA and 10 mM Tris, pH 8.1) and twice with wash buffer 4 (50 mM Tris, pH 7.4, and 50 mM NaCl). Proteins were eluted with 60 µl 1x Laemmli sample buffer (BioRad, 1610747) at 95**°**C.

### Immunoprecipitation

1.5×10^6^ HEK293T cells were plated on 60mm dishes and co-transfected the next day with 2µg of vectors encoding the GFP-tagged protein of interest and 2µg of a Cx36 expression vector. 24h after transfection cells were removed from the dish and centrifuged in phosphate buffered saline (pH 7.4) at 5000g for 5 min. The cell pellet was homogenized in immunoprecipitation buffer 250mM NaCl, 2m EDTA, 1% Tx-100, 50mM TrisHCL (pH 7.5) supplemented with protease inhibitor tablets (Roche). The homogenate was sonicated 3x for 30 sec and incubated on ice for 30 min. Afterwards homogenates were centrifuged for 10 min at 10,000g. 20µl of GFP-Trap agarose (Proteintech (Chromotek), was applied to the supernatant and incubated overnight on a rotating platform at 4°C. At the next day samples were centrifuged were centrifuged for 5 min at 2,500 g and washed three times in IP buffer. Proteins were eluted in 60 µl 1x Laemmli buffer (BioRad, 1610747) for 5 min at 95 °C.

### Western blot

Protein samples were separated via SDS-PAGE in a 10 % polyacrylamide gel at 160V. Afterwards proteins were transferred onto nitrocellulose membranes using the Trans-Blot Turbo transfer system (BioRad, 1704270, 1704159). Nitrocellulose membranes were blocked at RT in blocking solution containing 5% milk powder in TBST (20mM Tris pH 7.5, 150mM NaCl, 0.2 % Tween) and incubated in the primary antibody (diluted in blocking solution, Cx36 1:500, ThermoFisher, 37-4600, RRID:AB_2533320; Tubulin 1:1000, Abnova, MAB12827; Grasp55 1:1000, Proteintech, 10598-1-AP; GFP, 1:1000, Cell Signaling Technology, 2956, RRID:AB_1196615; V5 1:1000, ThermoFisher, R960-25, RRID:AB_2556564; Syntaxin5 1:5000, Synaptic Systems, 110 053; Sec24A 1:500, ThermoFisher, PA5-66043; Sec24B 1:10000, Bethyl Laboratories, A304-876A, RRID:AB_2621071; Ergic3 1:5000, Abcam, ab129179, RRID:AB_11141068; Sec24C 1:500, ThermoFisher, PA5-59101; GOPC: 1:1000, ThermoFisher, PA5-110897, RRID:AB_2856308) overnight at 4°C on a rotating platform. At the next day blots were washed 3x with TBST for 10 min and incubated with HRP conjugated secondary antibodies (goat anti mouse HRP, 1:1000, 34130 and goat anti rabbit, 1:1000, 34160, Thermofisher) for 1h. Nitrocellulose membranes were washed 3x with TBST and incubated for 1 min in ECL substrates (Thermofisher, 32106) for chemiluminescence detection.

### Imaging

Confocal scans were acquired with a Zeiss LSM800 airy scan microscope using a 60x oil objective (NA 1.3) and the airy scan function. Images were acquired as stacks consisting of 6-15 slices and recommended spacing of the airy scan mode (0.1-0.2 micron).

### Image analysis

Image analysis in this study was carried out in FIJI. For the quantification of PLA signals a rectangular region if interest (ROI) covering an area of 16.5µm^2^ was placed into perinuclear PLA signal. The mean intensity of PLA signal in this region was determined with the “Measure” function in FIJI. Colocalization was determined as previously described in Tetenborg et al., (19) using the colocalization highlighter plugin.

### Statistical analysis

Data sets acquired in this study were analyzed with GraphPad Prism 8. Data are shown as mean ± SEM. Normality was tested using the Anderson-Darling and the D’Agostino-Pearson test. Significance was tested using the two-tailed t-test or the two-tailed Mann-Whitney U test. For multiple comparisons a one way anova was performed.

## Results

### Functional ER export of Cx36 requires the C-terminal “YV” motif

We have previously shown that Cx36 mutants that lack the final four amino acids which constitute the PDZ binding motif (PBM) exhibit a trafficking defect that causes the connexin to accumulate in the ER. As a result of this retention, Cx36 channels inside the ER are allowed to dock prematurely leading to the formation of gap junction like aggregates that reshape the ER membrane into concentric whorls (Figure 1A). These structures can be found in the cytosol (illustrated in the cartoon in yellow, figure 1A) and around the nucleus (illustrated in magenta, figure 1A) and can be recognized via their characteristic toroid shape (whorls) in immunofluorescent imaging (19). In HEK293T cells transiently expressing wild type Cx36, the connexin forms perinuclear structures that colocalized with the SNARE proteins Syntaxin5 and Sec22, localizing at the ER-Golgi interface (Figure 1A). The association of Cx36 with both of these proteins implies, that the overexpressed connexin is trafficked via components of the secretory machinery present in HEK293T cells. Interestingly, although we observed functional gap junction formation, we found that Cx36 did not colocalize with endogenous ZO-1 in transfected HEK293T cells, suggesting that trafficking of connexons in the early secretory pathway is independent of ZO-1 in this context (Figure 1A).

**Figure 1.**
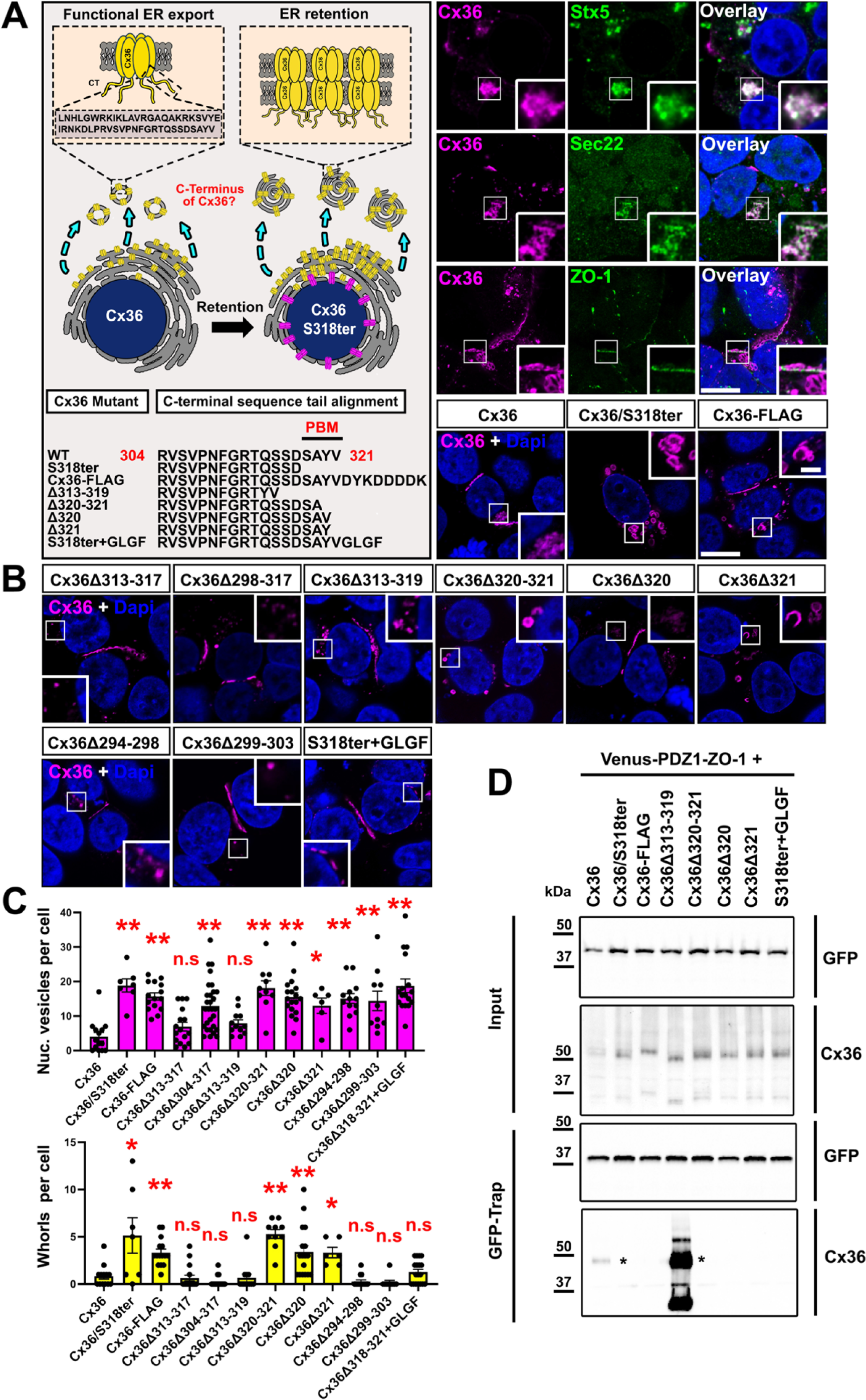
The C-terminal “YV motif” of Cx36 is necessary for ER export. **(A)** Left: Cartoon, illustrating the ER retention mechanism in C-terminal Cx36 mutants. Right: Colocalization of Cx36 with Stx5 and Sec22 in perinuclear regions. Cx36 in transfected HEK293T cells does not associate with endogenous ZO-1. **(B, C)** Insertions or deletions of amino acids at the very C-terminus result in ER retention. Deletions of amino acids N-terminal from the YV motif do not lead to ER retention and whorl formation. Scale in A and B: 10 µm. Values are shown as mean ± standard error of mean. Significance was tested using a one way anova. Cx36 mutants in which the very C-terminus was modified (mutation surrounding the YV sequence) show a significant increase in whorls and nucleus associated vesicles. P<0.01. **(D)** GFP trap pull-down of PDZ1 (ZO-1) co-expressed with different Cx36 mutants. Asterisk marks the Cx36 dimer band.

In the course of this study we used the PBM-deficient Cx36 whorl phenotype as a marker for ER retention to identify factors that control the functional ER export of Cx36. In a first set of experiments we asked whether the PBM alone, certain amino acids within the motif, or additional residues in the C-terminal tail mediate the ER export of Cx36. To identify a minimal motif we generated several Cx36 mutants that lack portions of the C-terminal tail and expressed these constructs in HEK293T cells. For each mutant, we determined the number of ER whorls in the cytosol (yellow bar graph) and vesicles that are surrounding the nucleus (magenta bar graph), as both of these structures are increasingly formed in HEK293T cells that express ER retained Cx36 mutants. Our mutagenesis screen revealed a significant increase in the number of ER whorls and nucleus associated puncta for Cx36 mutants that affected the very C-terminus comprising the amino acids tyrosine (Y) and valine (V) (Figure 1B and C). Attaching a FLAG tag to the C-terminus caused similar trafficking defects, suggesting that a functional carboxyl group of the valine residue is part of the export signal. This is consistent with the trafficking defects that have been described for the Cx36-EGFP construct (15, 16). When we deleted additional amino acids upstream from the YV motif and PDZ binding domain or directly within the C-terminal tail (Cx36Δ294-298, Cx36Δ299-303) we found no signs of ER retention. When we replaced the PDZ binding motif of Cx36 with an artificial sequence (GLGF) that matches the criteria for a class II PDZ binding motif (Φ-X-Φ-COOH), we observed an increase in the number of nucleus associated puncta but no signs of ER whorl formation.

Our mutagenesis screen demonstrated, that deletion of tyrosine 320 or valine 321 is sufficient to reproduce the ER export deficit we described for the S318/ter mutant. At this point, however, it is unclear if the ER export defect arises from the loss of a separate export signal or from the disruption of PDZ domain mediated interactions. To address this issue, we tested the ability of each PBM mutant to bind the PDZ1 domain of ZO-1 (Figure 1D). We co-expressed a venus-tagged PDZ1 domain with each Cx36 mutant in HEK293T and performed a GFP trap pulldown. Of the constructs tested, those which showed the whorl phenotype were unable to bind, and only wild type Cx36 and the Cx36Δ313-319 bound PDZ1. Additionally, we found that the Cx36Δ313-319 mutant showed a massive increase in PDZ binding, although it is lacking the first two amino acids of the PBM. This drastic effect is likely due to the repositioning of the YV motif next to the upstream sequence RT creating a high affinity PDZ ligand with following sequence: RTYV. Taken together, we find that Cx36 mutant that showed ER retention (whorls) also failed to bind PDZ1, raising the question whether functional ER export requires the entire PBM “SAYV” or the “YV” motif.

### Functional ER export of Cx36 requires the cargo receptors Sec24A and B

Our data have shown that an intact C-terminus of Cx36 determines the functional ER export of Cx36. The release of newly synthesized proteins from the ER in eukaryotic cells is regulated by the COPII complex. This conserved release machinery requires the assembly of cytosolic coat proteins that deform the ER membrane at ER exit sites into secretory vesicles containing the cargo protein (22). Within the COPII complex cargo recognition is controlled by Sec24 isoforms, a family of adapter molecules that connect the cargo protein to the COPII coat via interactions with certain export signals in its cytoplasmic domains (23). As the COPII complex functions as an almost universal export route for most proteins, we reasoned that the trafficking deficient Cx36 mutants may fail to interact with the Sec24 cargo receptors preventing them from entering the COPII vesicle and resulting in ER retention. To address this hypothesis, we first tested if inhibition of the COPII complex reproduces the phenotype we observed for trafficking deficient Cx36 mutants. The assembly of the COPII coat is initiated by the small G protein Sar1. We co-expressed the dominant negative Sar1 (T39N)-GFP mutant, which serves as a competitive inhibitor of endogenous Sar1 (24), with wild type Cx36 to block COPII complex formation and analyzed the intracellular distribution of the connexin. Sar1(T39N)-GFP co-transfection caused an accumulation of Cx36 into dense perinuclear vesicles (magnified inset) and the formation of smaller aggregates surrounding the nucleus (white arrows) (Figure 2A). Especially the nucleus associated puncta (white arrows) were reminiscent of the trafficking defect we have described for Cx36 mutants lacking the PDZ binding motif. Interestingly, we observed that co-transfection of the COPII coat protein Sec23 disrupted the transport of wild type Cx36 in a similar way as removal of the PDZ binding motif. Sec23-GFP transfected cells formed the characteristic connexin whorls (magnified inset) and smaller aggregates surrounding the nucleus (white arrows) (Figure 2A). Although Sec23-GFP doesn’t function as a COPII inhibitor, it is likely that the overexpressed coat protein depleted binding sites for the endogenous Sec23 within the COPII complex and thereby prevented the release of newly synthesized Cx36 channels. When we co-expressed Sec61B-GFP, an ER membrane protein that does not belong to the COPII complex, we did not see any signs of ER retention, confirming that ER export of Cx36 is COPII dependent.

**Figure 2.**
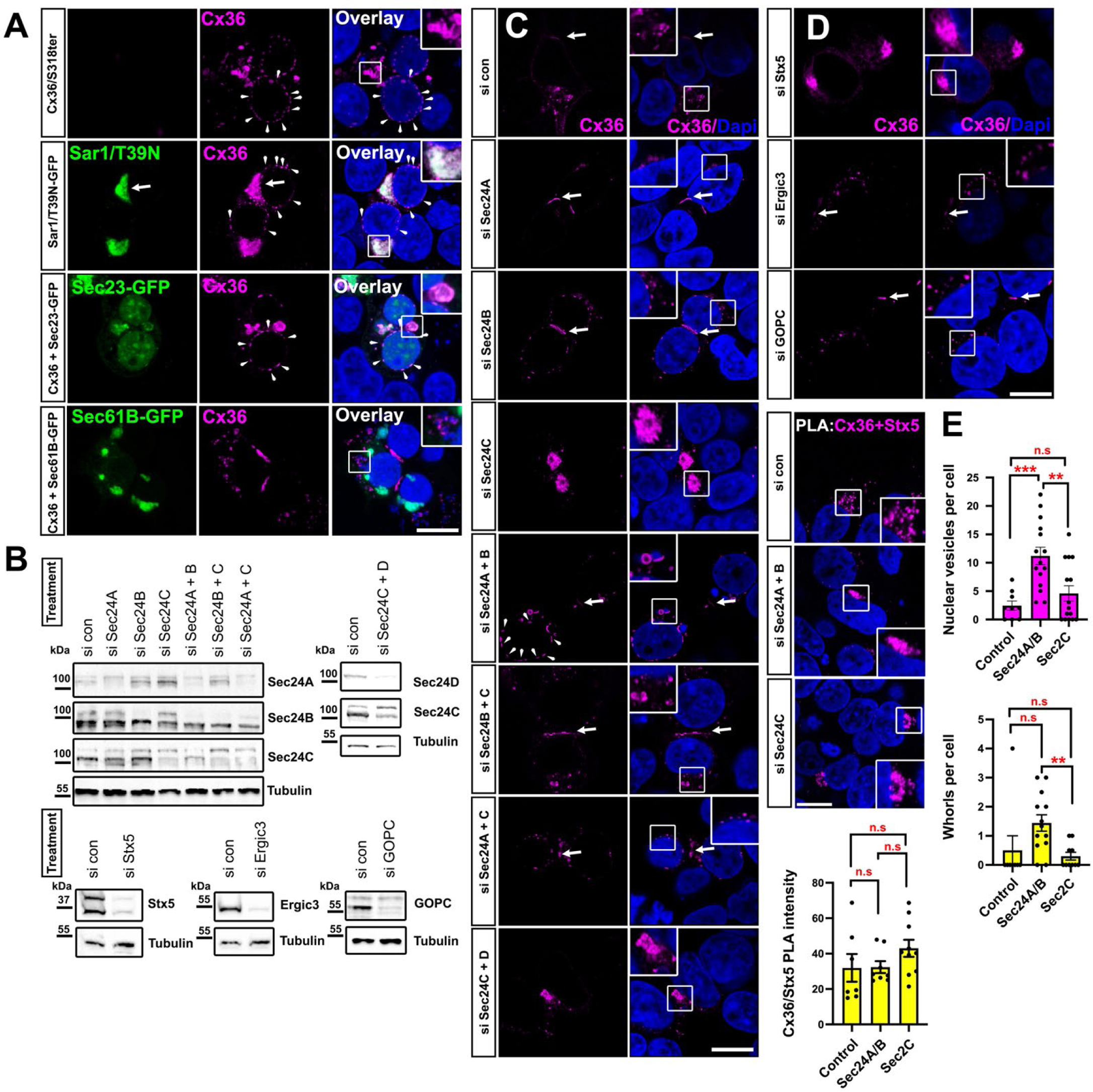
Cx36 requires COPII cargo receptors for ER export. **(A)** Overexpression of Sar1T39N-GFP and Sec23A-GFP lead to accumulation of Cx36 around the nucleus as previously described for the Cx36/S318ter mutant. **(B)** Western blots confirming siRNA mediated depletion of secretory pathway proteins. **(C)** Confocal scans of Cx36 in siRNA treated cells. Knock-down of specific Sec24 isoforms has distinct effects on the distribution of Cx36. Combined knock-down of Sec24A and B reproduces the trafficking defect observed for the Cx36/S318ter. Knock-down of Sec24C, on the contrary, leads to accumulation of Cx36 in perinuclear structures. **(D)** Knock-down of ER-Golgi trafficking proteins Ergic3 and GOPC have no effect on Cx36 distribution, while knock-down of Syntaxin5 leads to accumulation of Cx36 in the perinuclear region. **(E)** Evaluation of different phenotypes observed in Sec24A/B or Sec24C depleted cells. Cx36/Stx5 PLA intensity was not significantly different. All p values were > 0.05. More ER whorls are formed in Sec24A/B depleted cells. Data are shown as mean ± standard error of mean. Significance was tested using a one way anova. Scale: 10µm.

Next, we sought to identify the exact cargo receptor isoforms that bind to Cx36 in the COPII complex. We targeted all four Sec24 variants using siRNA and tested how silencing of each isoform influenced the intracellular distribution of Cx36. To validate the successful knock-down we tested the Sec24 protein level in cell lysates from each transfection (Figure 2B). Knock-down of Sec24 A and B showed little to no effects on the distribution of Cx36. Only combined depletion of both isoforms led to the formation of the characteristic Cx36 whorls (magnified insets) and aggregates surrounding the nucleus (Figure 2C), resembling the trafficking defect we have described for the Cx36S318Ter (Figure 1B). Silencing of Sec24C or double knockdown of Sec24C and D, on the contrary, concentrated Cx36 into dense perinuclear structures.

Besides the Sec24 isoforms, we tested the impact of additional proteins that have been linked to intracellular trafficking events (Figure 2D). Among them was Ergic3, a membrane protein that has been shown to control the ER to Golgi transition of Connexin43 and innexins (invertebrate GJ forming proteins (25)), suggesting a function as a universal export factor for metazoan gap junction proteins (26). In our hands, however, depletion of Ergic3 did not cause any obvious signs of ER retention. Next, we targeted the Golgi specific PDZ protein GOPC (Golgi-associated PDZ and coiled-coil motif-containing protein). This protein appeared as an interesting candidate due the PDZ domain and role in the trafficking of claudin1 and 2, which carry C-terminal YV motifs identical to Cx36 (27). Similar to the results for Ergic3, we did not observe any ER retention effects for Cx36 when we silenced GOPC. As a final candidate on our list we targeted Syntaxin5 (Stx5) a SNARE protein that fuses the membranes of ER export vesicles and the cis Golgi to allow the uptake of secreted cargos into the Golgi apparatus (28). Interestingly, we found that depletion of Stx5 caused the formation of dense perinuclear clusters containing Cx36. As Stx5 depletion per se would not prevent the release of Cx36 from the ER, it is likely that these clusters represent secreted channels that accumulate around the Golgi apparatus.

We observed that silencing of Sec24A and B or Sec24C resulted in quite distinct intracellular distributions of Cx36. While depletion of Sec24A/B reproduced the ER retention phenotype for Cx36, knockdown of Sec24C, on the contrary, led to the formation of dense perinuclear clusters resembling the Golgi apparatus. We further evaluated these phenotypes and performed a proximity ligation assay to detect Cx36/Stx5 interactions. This strategy allowed us to label Cx36 channels that have transitioned into the Golgi apparatus since Stx5 is largely confined to the cis Golgi. In comparison to control siRNA treated cells, however, this assay only detected a subtle insignificant increase in Cx36/Stx5 in Sec24C depleted cells. The number of ER whorls in Sec24C depleted cells was significantly lower in comparison to Sec24A/B depleted cells, reflecting the lack of an ER retention phenotype when Sec24C was knocked down. In comparison to control siRNA treated cells, Sec24A/B depletion substantially increased the number of ER whorls (yellow bar graph) and vesicles surrounding the nucleus (Figure 2E). While the increase in the frequency of nucleus associated vesicles in Sec24A/B depleted cells was highly significant compared to the control condition, the number of cytosolic whorls (yellow bar graph), however, was not. Though it appears, that Sec24A/B depletion reproduced only one of the two ER retention phenotypes, one has to consider that in comparison to simple overexpression experiments (Figure 1) a lower amount of the Cx36 expression vector (333 ng) had to be co-transfected with Sec24 siRNAs to avoid cytotoxic effects. Since whorl formation heavily depends on the amount of Cx36 localizing in the ER, it is likely that our assessment of ER whorls in Sec24A/B depleted cells is an underestimate of the actual ER the retention effect.

To understand how Cx36 interacts with the COPII cargo receptors, we first analyzed the colocalization of the connexin with Sec24 A, B and C. Each of these receptors colocalized with Cx36 in perinuclear regions (Figure 3A). Consistent with the knock down experiments, we observed a strong association of Cx36 with Sec24A and B, with around 60% of perinuclear Cx36 colocalizing with the two cargo receptors (Figure 3B). A lower degree of colocalization was detectable for Sec24C. We next asked whether the cargo receptors Sec24B and C directly bind to Cx36 and co-transfected GFP-tagged Sec24 constructs with Cx36 into HEK293T cells to perform a pull-down assay (Figure 3B). As a positive control we transfected the venus tagged PDZ1 domain of ZO-1 which showed substantial binding to Cx36. We observed no association with the connexin when the pull-down was performed with the Sec24B or C, suggesting that Cx36 does not directly interact with these cargo receptors. Another possibility to explain the lack of a detectable interaction is that Sec24 isoforms bind Cx36 with low affinity making co-precipitation experiments difficult. To bypass this issue and test a molecular association of Sec24B and Cx36 we performed a proximity ligation assay using the duo link PLA system. We transfected Cx36 into HEK293T cells and tested the PLA reactivity targeting Cx36 and endogenous Sec24B. Consistent with the double labeling experiments we observed strong PLA reactivity in perinuclear regions likely reflecting ER exit sites (Figure 3C). A clear reduction in PLA labeling of these perinuclear regions was observed when the assay was performed with Cx36/S318Ter mutants lacking a functional C-terminus. Based on this observation we conclude that a molecular association with Sec24B involving a functional C-terminal tip of Cx36 is necessary to incorporate the connexin into COPII vesicles and transport it out of the ER.

**Figure 3.**
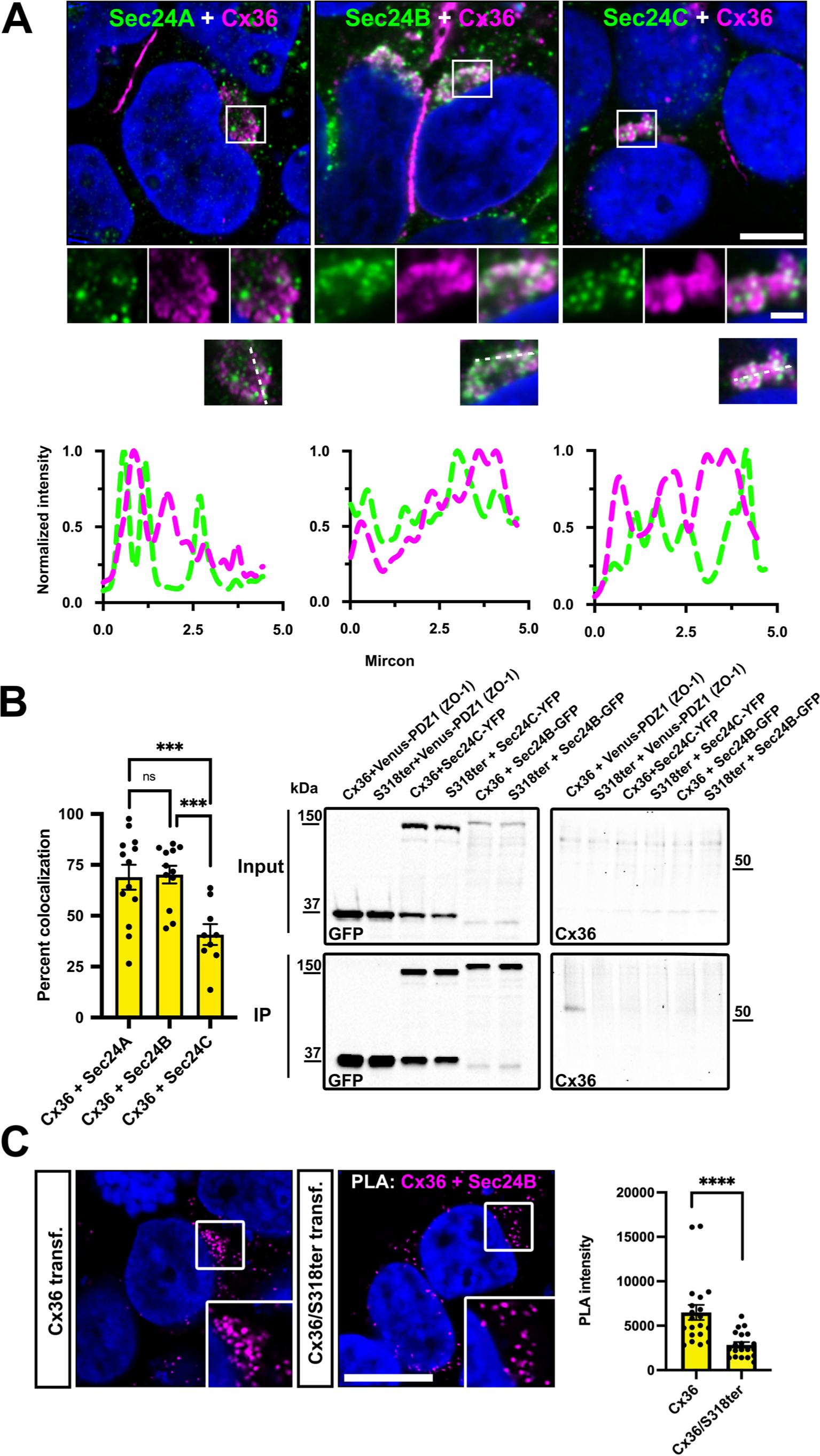
Cx36 associates with COPII cargo receptors. **(A)** Cx36 shows partial colocalization with the three different cargo receptors Sec24A, Sec24B and Sec24C. Line scans depict the overlap of intensity maxima for a selection region of interest. **(B)** Colocalization analysis illustrates the degree of colocalization for Cx36 and different COPII cargo receptors. The difference in Cx36 colocalization for each Sec24A and B was significant in comparison to Sec24C. Sec24A vs. Sec24C, P=0.0032. Sec24B vs. Sec24C, P= 0.0024. Values are shown as mean ± standard error of mean. Significance was tested using a one way anova. Immunoprecipitations fail to detect binding interactions of Cx36 and the COPII cargo receptors Sec24B and Sec24C. Venus-PDZ1-ZO-1 served as a positive control and showed a robust interaction with Cx36. **(C)** Proximity ligation assay was used to assess the molecular association of Cx36 and Sec24B. The Cx36 and Sec24 PLA reaction results in compact perinuclear signals that are significantly weaker in intensity when the assay is performed with Cx36/S318ter. Scale: 10 µm.

### Cx36 interacts with the Golgi stacking protein Grasp55

To identify additional proteins acting within the secretory pathway to control Cx36 trafficking, we used a TurboID screening approach. TurboID is an engineered, bacterial enzyme that promiscuously biotinylates lysine residues on proteins within an ∼10nm radius of localization, subsequently allowing for easy isolation of the biotinylated proteins (29). We first tried engineering TurboID directly onto Cx36, but found that insertion of various TurboID variants into Cx36 interfered with the functional transport of gap junction channels, irrespectively of the insertion site. We therefore turned to the recently developed BLITZ technique, in which a destabilized GFP-targeting nanobody carries TurboID to subcellular locations where proteins are tagged with GFP (19, 21) (Figure 4A). Given that we found that disrupting the terminal “YV” motif resulted in ER export defects (Figures 1-3), we co-expressed wildtype Cx36 and Cx36-EGFP in HEK293T cells and found that the tagged Connexin did not display the hallmarks of ER retention (Figure 4A). The Cx36-EGFP construct is presumably being rescued by the wildtype Cx36 that is co-expressed, as has been observed *in vivo* in transgenic mice (15, 16). We additionally co-expressed the destabilized GFP-targeting nanobody that carries TurboID (V5-TurboID-dGBP) and found that it colocalizes with Cx36-EGFP and results in substantial biotinylation at the plasma membrane (Figure 4A arrows) and intracellular vesicles (Figure 4A small arrow heads). This control experiment confirms that the TurboID construct is correctly targeting Cx36 without compromising the transport of gap junction channels. We isolated biotinylated proteins from Cx36-EGFP/Cx36/ V5-TurboID-dGBP co-transfected HEK293T cells, and as control, cells expressing only V5-TurboID-dGBP, and analyzed the samples via mass spectrometry. In Cx36-EGFP/Cx36 co-transfected samples we identified several secretory transport proteins including Sec24A/B, synergin gamma, and the Golgi reassembly stacking protein 55 (Grasp55) (Figure 4A and B). Grasp55 was of particular interest for us given its function in unconventional protein secretion (30) and its two PDZ domains mediating interactions with proteins bearing C-terminal valines such as CD8 (31). We first tested if Cx36 and Grasp55 interact and co-expressed GFP-tagged Grasp55 with Cx36. We isolated overexpressed Grasp55 from co-transfected HEK293T cells using GFP trap and co-precipitated substantial amounts of Cx36. When the pull-down was performed with a Grasp55 construct lacking the two PDZ domains or with a truncated mutant of Cx36 missing the PBM we observed no binding, indicating that Grasp55 interacts with Cx36 via its PDZ domain (Figure 4C). We further observed that transfected Cx36 colocalized with endogenous Grasp55 in perinuclear structures resembling the Golgi apparatus (Figure 4D). To demonstrate that Cx36 also interacts with endogenous Grasp55 in HEK293 we performed a proximity ligation assay and measured the intensity of perinuclear PLA signals in Cx36 transfected cells (Figure 4E). In comparison to wild type Cx36 transfected conditions the Cx36S318ter mutant showed significantly weaker Cx36/Grasp55 PLA reactivity, suggesting that Cx36 binds to endogenous Grasp55 in a PDZ dependent manner. Besides Grasp55 we identified the protein tyrosine phosphatase non-receptor type 13 (PTPN13) in our BioID screen as an additional interactor of Cx36. PTPN13 consists of multiple of PDZ domains and functions as a tumor suppressor (32). We have previously shown that perch Cx35, which has an identical PBM to Cx36, bound to the second PDZ domain of PTPN13 in a micro array screen (33). Consistent with this finding, we observed extensive colocalization of transiently expressed Cx36 and endogenous PTPN13 at gap junctions in transfected HEK293T cells (Figure 4F). We excluded PTPN13 as an additional secretory factor for Cx36 because the protein was mostly confined to gap junctions but rarely associated with intracellular vesicles. Thus, the only PDZ protein we identified in our BioID screen that could potentially affect trafficking of Cx36 in early secretory compartments is Grasp55.

**Figure 4.**
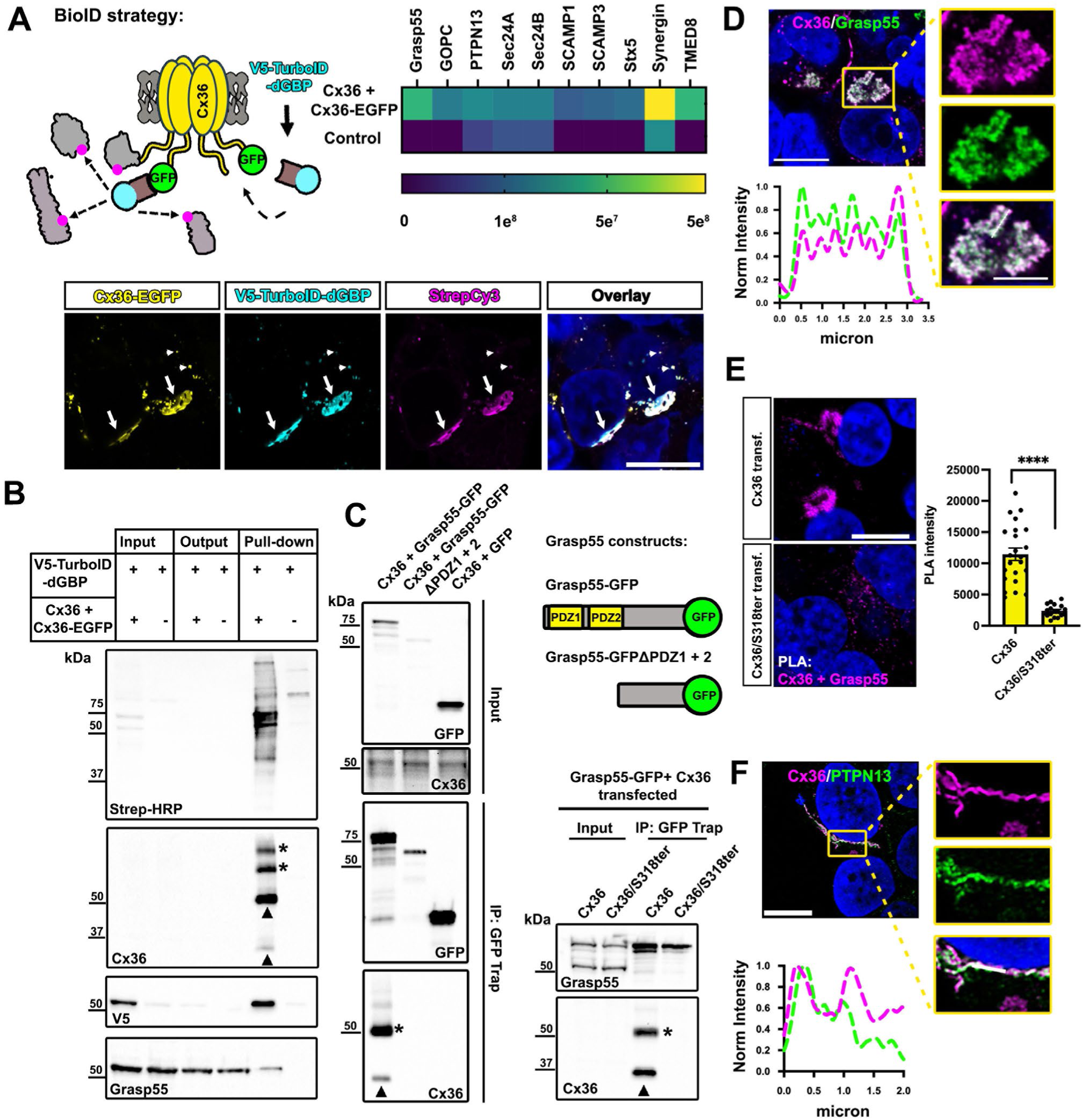
Cx36 and Grasp55 interact via PDZ/PBM binding and colocalize in the Golgi. **(A)** GFP directed proximity biotinylation was used to identify proteins regulating the transport of Cx36 in the secretory pathway. Immunostaining confirms correct targeting of V5-TurboID-dGBP and localized biotinylation. Heatmap summarizes the abundance of hits in the BioID screen. Among several secretory pathway proteins we identified Grasp55 in Cx36/Cx36-EGFP transfected conditions. **(B)** Streptavidin-affinity-captured proteins were detected on western blots using Streptavidin-HRP, anti-Cx36, anti-V5 and anti-Grasp55. **(C)** Immunoprecipitation using GFP trap demonstrates a PDZ dependent binding interaction between Cx36 and Grasp55-GFP. A Grasp55 variant lacking the PDZ binding motif fails to interact with Cx36. A Cx36 mutant lacking the PDZ binding motif does not co-precipitate with full length Grasp55-GFP. **(D)** Confocal scan of Cx36 and Grasp55 colocalizing in perinuclear structures. Line scan illustrates the overlap of intensity maxima. **(E)** Proximity ligation assay confirms a molecular association of Cx36 and Grasp55 that is diminished when the Cx36/S318ter mutant (lacking PDZ binding motif) is transfected. **(F)** Colocalization of PTPN13 and Cx36 at gap junctions in HEK293T cells. Difference in PLA labeling was highly significant. Scale: 10 µm.

### Grasp55 concentrates Cx36 in the Golgi

To further characterize the interaction between Cx36 and Grasp55, we first determined the subcellular compartment in which these proteins associate. In contrast to previous reports (34), we identified Grasp55 in the cis-Golgi as indicated by the colocalization with the Golgi matrix protein GM130. By contrast, the trans Golgi marker golgin97 showed no signs of association with Grasp55, suggesting that the scaffold protein is mainly restricted to the cis Golgi (Figure 5A). Additionally, we observed some association of Sec24B and Grasp55, which is consistent with previous reports (20). Co-expression of Cx36 and Cx36-EGFP combined with double labeling of organelle markers and Grasp55 further confirmed that Cx36 and Grasp55 colocalize in the cis Golgi, colocalized with GM130 (Figure 5B). In the corresponding intensity scans a clear overlap for the maxima of all three channels can be seen. To investigate how Grasp55 impacts the vesicular transport of Cx36, we expressed the connexin in Grasp55 deficient HEK293T cells previously generated by Liu et al., (20) and quantified gap junction size, frequency and Cx36-containing ER whorls as an indicator of ER retention (Figure 5C). Surprisingly, we found that Grasp55 deletion influenced none of these parameters. Compared to Cx36 transfected control HEK293T cells, Grasp55 KO cells showed no changes in gap junction size, frequency or ER whorls, indicating that Grasp55 is dispensable for the functional ER export of Cx36 and gap junction formation. While deletion of Grasp55 caused no apparent transport deficits, overexpression of a Grasp55-EGFP construct led to retention of Cx36 in perinuclear structures resembling the Golgi apparatus. This retention effect was prevented when the Grasp55 G2A mutant was transfected (Figure 5D). The G2A mutant unlike wild type Grasp55 is not myristoylated at the N-terminus and fails to localize to the Golgi apparatus (35), suggesting that the retention effect of Grasp55 is caused by an interaction with the Golgi membrane. To test if Grasp55 affects the transition of Cx36 into the cis-Golgi we compared Cx36/Stx5 PLA reactivity between wild type and Grasp55 knock out cells. We observed no significant reduction in Cx36/Stx5 reactivity indicating that the Grasp55 deficiency does not affect the transition of Cx36 into the Golgi apparatus (Figure 5E). Thus, we conclude that Grasp55 is dispensable for the ER export of Cx36 but could be necessary to concentrate the connexin in the Golgi apparatus.

**Figure 5.**
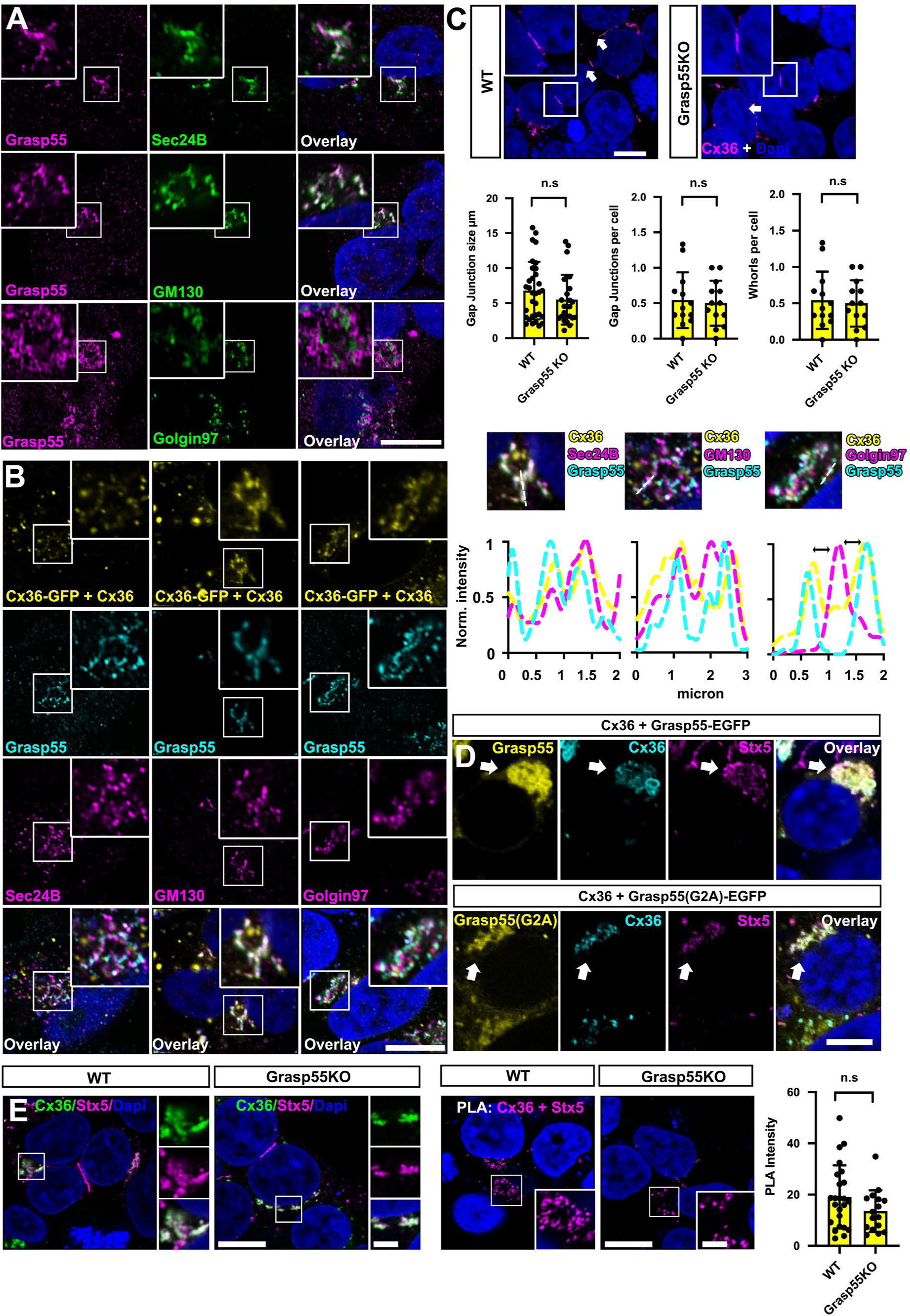
Grasp55 localizes to the cis Golgi and where it can retain Cx36. **(A)** Grasp55 resides in the Cis-Golgi and colocalizes with GM130 but not with golgin97. colocalize in the cis-Golgi. No colocalization can be detected in the trans-Golgi or ER exit sites. **(B)** Grasp55 and Cx36 (visualized as Cx36-EGFP/Cx36 heteromers. Detection of Cx36-EGFP via GFP allows immunolabelling of three channels, since mouse and rabbit antibodies were already used for organelle markers. Coexpression of native Cx36 is necessary to maintain functional PDZ interactions) **(C)** Knockout of Grasp55 does not compromise gap junction formation and Cx36 trafficking. Differences between conditions were not significant, P>0.05. Significance was determined using a Mann-Whitney U test. **(D)** Overexpression of Grasp55 leads to retention of Cx36 in perinuclear structures. **(E)** PLA labeling of Stx5 and Cx36. Knockout of Grasp55 does not affect the transition of Cx36 into the Golgi. Difference in Cx36/Stx5 PLA intensity was not significant between wild type HEK294T cells and Grasp55 KO cells. P>0.05. Significance was determined using a Mann-Whitney U test. Scale: 10 µm.

## Discussion

In the present study we have investigated the molecular mechanisms that guide the transport of Cx36 in the early secretory pathway. By combining RNAi and GFP dependent proximity biotinylation assays we identified functional interactions of Cx36 and two distinct protein complexes mediating secretory trafficking: The cargo receptors Sec24A and B of the COPII complex and the Golgi stacking protein Grasp55 in the cis-Golgi. The interaction of Cx36 with the COPII complex involves the C-terminal amino acids tyrosine and valine while binding of Grasp55 requires the PDZ binding motif.

The function of a C-terminal valine serving as an ER export signal is not a unique feature that only applies to Cx36. Around 10 % of human type I membrane proteins display a valine at the extreme C-terminus (36) and several studies have characterized its respective function as an ER export signal in expression systems. The deletion of this residue in CD8 (37), Kitl (kit ligand) (38) and HLA-F (human leucocyte antigen F) (39) prevents the functional ER export as we have described for Cx36, suggesting that a more general mechanism is involved. This is not surprising given that the COPII complex accounts for 2/3 of all proteins that have to be exported from the ER. Although the combined depletion of Sec24A and B replicated the trafficking defect we have described for the Cx36S318ter mutant, we weren‘t able to demonstrate a direct binding interaction between Cx36 and these two cargo receptors, which is surprising since our co-precipitation assay was sensitive enough to capture low affinity complexes that are formed by Cx36 and the first PDZ domain of ZO-1 (40). It is possible that the interaction of Cx36 and Sec24A/B requires a yet unidentified adapter protein that binds to both proteins simultaneously thereby linking Cx36 indirectly to the COPII coat. Indeed Sec24 proteins are known to recruit additional cargo receptors such as Erv14 which have unique binding preferences that diversify the cargos that are sorted into COPII vesicles (41). Moreover, we can‘t exclude that the affinity between Cx36 and Sec24A/B is extremely low, since the export signal consists of two amino acids which would create a short binding interface. Despite the lack of a detectable interaction in our IP experiments, we observed strong PLA labeling for Cx36 and endogenous Sec24B that was significantly reduced in intensity when the Cx36S318ter mutant was expressed. Based on this observation, we conclude that Cx36S318ter mutant fails to bind Sec24A/B (directly or indirectly) preventing it from entering the COPII vesicle. As a result, this mutant is retained in the ER giving rise to whorl formation as described previously.

A previous study by Yin et al. (42) has shown that the tight junction protein claudin1 binds to Sec24C via the C-terminal YV motif, a short sequence that is found in many claudin isoforms and Cx36. Functional ER export of claudin1 can be blocked entirely by depletion of Sec24C. Although Cx36 exhibits a very similar export signal, we found no evidence suggesting that Sec24C functions as an exclusive export factor for Cx36. Although we did not assess direct binding interactions between Cx36 and the different Sec24 isoforms, we found that silencing of Sec24A/B or C had quite distinct effects on the localization of the connexin. While combined depletion of Sec24A and B resulted in obvious ER retention, as indicated by the formation of nucleus associated vesicles, silencing of Sec24C on the contrary often led to the formation of dense perinuclear clusters. One possible interpretation of these findings is that Sec24C is necessary for Cx36 to bypass the Golgi, which would explain why depletion of this cargo receptor enhanced the transition of Cx36 into the Golgi apparatus. Indeed, Golgi bypass mechanisms have been observed in neurons and play a role in targeting NMDA receptors into dendritic compartments (43). Moreover, Sec24C has been shown to mediate a Golgi bypass mechanism for ABC transporters in *Arabidopsis thaliana* (44). Thus, considering the diversity of neuronal compartments in which electrical synapses are formed (16, 45), it seems feasible that discriminatory transport mechanisms are required to direct Cx36 to the correct location. An interaction with different cargo receptor variants of the COPII complex could represent one of these mechanisms. However, further *in vivo* studies are necessary to elucidate the role of Sec24 isoforms in synaptic targeting of Cx36.

Grasp55 has been shown to be a versatile protein, and was initially described as a second Golgi stacking protein forming oligomers that link the cisternae to maintain the characteristic structure of the Golgi apparatus (35). More recent studies have focused on its function as a secretory factor (46). Gee et al., reported that Grasp55 mediates the unconventional secretion of the cystic fibrosis transmembrane conductance regulator (CTFR) via a Golgi-independent pathway. This mechanism is activated by ER stress and rescues the surface expression of the disease-causing ΔF508 mutant (30). We cannot entirely exclude that Grasp55 supports the transport of Cx36 in a similar way, especially since we detected Cx36/Grasp55 PLA signals at gap junctions and outside the Golgi apparatus. However, most of the PLA reactivity was localized to the cis Golgi, which argues for a more spatially confined function. Moreover, we found no signs of ER retention nor any impairments in gap junction formation in Grasp55 KO cells, suggesting that Sec24A/B but not Grasp55 is the main force driving the ER export of Cx36. Our co-transfections further demonstrate that heterologously expressed Grasp55, unlike the G2A mutant, causes Cx36 to accumulate in the Golgi. This retention effect is caused by a simultaneous interaction of Grasp55 with the Golgi membrane and Cx36, tying the connexin to the Golgi. This clearly raises the question of how such a mechanism is supposed to support the transport of a synaptic protein? A recent study by Pothukuchi et al. (34) has shown that Grasp55 regulates the localization of glycosphingolipid biosynthesis enzymes inside the Golgi. The correct targeting of these enzymes is determined by an equilibrium of retrograde carriers and the stabilizing function of Grasp55. In the absence of Grasp55, glycosphingolipid biosynthesis enzymes are increasingly sorted into COPI vesicles, preventing their compartmentalization in the trans Golgi (34). This observation suggests that Grasp55 is necessary to stabilize the localization these enzymes in order to prevent their transition into retrograde carriers. The membrane proteins CD8 and the Fz4 receptor have been shown to interact with Grasp55 and 65, respectively. Both receptors display C-terminal valine residues and their interaction with either Grasp55 or Grasp65 accelerates the transport through the Golgi (31). Although it is unknown how exactly Grasp proteins impact the transport of their binding partners, it was proposed that interactions with the PBM neutralize retrograde trafficking signals and thereby facilitate anterograde transport mechanisms. A similar mechanism might apply to Cx36, which would make its transport more efficient.

In summary, our study provides a first detailed description of the initial events that determine the intracellular transport of Cx36 and provide a mechanistic explanation for trafficking defects that have been described in previous publications (15, 16, 19).

## Acknowledgements

We would like to thank Ya-Ping Lin and Nikki Brantley for the excellent technical assistance. We’d like to thank Li Li from the Clinical and Translational Proteomics Service Center, UTHealth, for mass spectrometry sample services.

## Data availability

All methods and data included in the manuscript will be made available upon request to Dr. Stephan Tetenborg or Dr. John ÒBrien.

---

**Figure 6.**
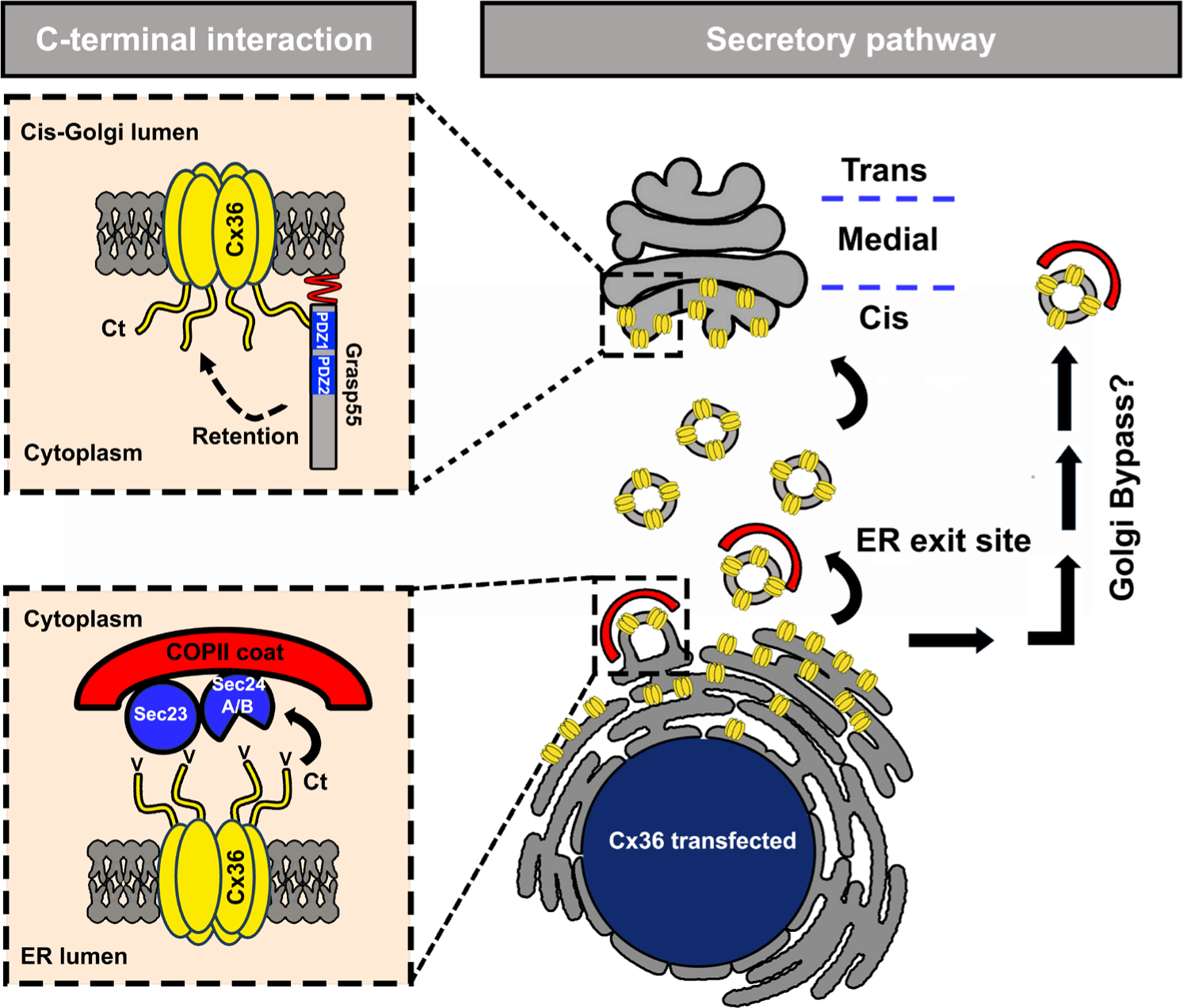
Summary model illustrating the transport mechanisms of Cx36 in the early secretory pathway. The C-terminus of Cx36 and Sec24A/B mediate the uptake of Cx36 into COPII vesicles. Once Cx36 transitioned into the cis-Golgi, it interacts with Grasp55 via its PDZ binding motif, resulting in Golgi retention. A pathway bypassing the Golgi may also provide a route for Cx36 to be transported to the plasma membrane.

